# Spatiotemporal analysis of *de novo* KSHV infection using Crispr/Cas9-based 3D live cell imaging at single episome resolution

**DOI:** 10.1101/2025.02.11.637615

**Authors:** Thomas Günther, Simon Weissmann, Martin V. Hamann, Henry Scheibel, Jens B. Bosse, Marion Ziegler, Adam Grundhoff

## Abstract

Kaposi Sarcoma-associated herpesvirus (KSHV) persists as a latent episome in infected cells. While the virus efficiently infects established cell lines and primary cells *in vitro*, the early events guiding establishment of latent infection and the dynamic interplay between viral episomes and host factors remain incompletely understood. Here, we describe the development and application of a CRISPR/Cas9-based 3D live cell imaging system capable of tracking single KSHV episomes in real-time. Our approach exploits the SunTag technology, wherein deactivated Cas9 (dCas9) molecules are fused to repetitive epitope arrays recognized by superfolder GFP-fused single-chain antibodies. By targeting these complexes to terminal repeat units of KSHV, we achieve high level signal amplification, allowing us not only to detect newly incoming viral genomes within the first hours of *de novo* infection, but also to follow their spatiotemporal trajectories through different stages of the viral lifecycle. Furthermore, to facilitate efficient generation of stable reporter cell lines, we adapted the transposon-based piggyBac system to combine all SunTag components into a single-vector targeting system (SunSeT).

Using these systems, we demonstrate the ability to observe both transient and stable interactions between KSHV episomes and key cellular regulators, including the variant polycomb-repressive complex 1 (vPRC) component KDM2B and the innate immune sensor IFI16. Furthermore, our platform allows detailed visualization of episodic changes in episome localization, abundance and distribution during *de novo* and long-term infection, providing critical insights into how viral genome positioning and dynamics correlate with host subnuclear environments.

Overall, our study introduces a robust and adaptable imaging platform to dissect the earliest events of KSHV infection. The ability to track viral episomes in living cells offers a powerful tool to advance our understanding of the spatial and temporal regulation of individual KSHV genomes, shedding light on fundamental mechanisms of herpesvirus latency and persistence.

## Introduction

Over the past decade, significant advancements have been made in the detection and visualization of DNA in living cells, providing crucial spatiotemporal insights into locus-specific DNA interactions with other DNA loci, RNA, and proteins (reviewed in ^1–3^). These methods enable single-cell and single-locus investigations, allowing for precise tracking of genomic dynamics, however, most techniques require genetic manipulation of the targeted locus.

During *de novo* infection, non-chromatinized and epigenetically naïve linear KSHV DNA is injected from the viral capsid through the nuclear pore into the nucleoplasm. In this early phase, the viral genome undergoes circularization and chromatinization, acquiring an orchestrated sequence of activating and repressive histone modifications. These modifications establish compatibility with the subsequent expression programs governing latency and lytic replication. Traditionally, investigations of this critical phase relied on bulk chromatin techniques, such as chromatin immunoprecipitation (ChIP), or imaging methods like fluorescence *in situ* hybridization (FISH) combined with immunofluorescence analysis (IFA). While these approaches have provided valuable insights, they lack the ability to capture dynamic interactions of viral episomes in living cells.

The introduction of DNA labeling technologies, including click chemistry and capsid labeling via fluorescent protein fusion to tegument proteins, has facilitated imaging of different stages of the herpesvirus life cycle. For example, herpes simplex virus 1 (HSV-1) can incorporate EdC and EdU into its genome during lytic replication^4^. However, such approaches face limitations: Click chemistry signals are lost upon latent DNA replication, EdU/EdC incorporation may be virus-specific, and cells frequently are fixed for the click reaction. Similarly, capsid protein labeling is primarily useful for studying viral entry and egress rather than nuclear interactions during latency. Alternative approaches, such as the ANCHOR system^5^ for visualizing viruses e.g. human cytomegalovirus DNA^6^ or imaging using lac operon arrays^7^ and similar technologies, offer powerful tools but are constrained by the requirement for recombinant or BAC-engineered viruses, making them less suitable for studying wild-type KSHV infections.

Robust techniques capable of specifically tracking the fate of individual KSHV episomes from nuclear entry through multiple cell divisions and lytic replication have been lacking. To address this methodological gap, we employed the previously reported SunTag system^8^, to enable live-cell tracking of KSHV episomes throughout the earliest stages of infection. Furthermore, we combined the lentivirus-based components of the original SunTag system into a transposon-based piggyBac integration vector^9,10^. Our single-vector targeting system (SunSeT) enables fast and robust generation of KSHV reporter cell-lines by transfection. This system allows for the visualization of transient and stable interactions between viral DNA and key cellular regulators, including the variant polycomb-repressive complex 1 (vPRC) component KDM2B and the innate immune sensor IFI16. By enabling real-time tracking of viral episomes, the SunTag system overcomes the limitations of previous approaches, providing novel insights into the spatial and temporal regulation of KSHV genomes within the host nucleus. This study introduces a powerful tool for dissecting the earliest events of KSHV infection, offering new avenues for exploring herpesvirus biology and the interplay between individual viral episomes and host cell processes.

## Results

### Establishment of the SunTag-based KSHV live-cell tracking system

To visualize and track *de novo* infecting KSHV episomes over extended periods during infection, we first established a stable SunTag expression cell line via lentiviral transduction.

Initially, we employed the original lentiviral SunTag expression plasmids, pHRdSV40-NLS-dCas9-24xGCN4_v4-NLS-P2A-BFP-dWPRE and pHR-scFv-GCN4-sfGFP-GB1-NLS-dWPRE, as described in the foundational SunTag publication by Tanenbaum and colleagues^8^. These constructs lack a selectable antibiotic resistance gene but contain blue and green fluorescent proteins, respectively. However, the lentiviral titers we obtained with the dCas9-24xGCN4-BFP construct were insufficient for generating stable cell lines, as determined by BFP fluorescence-activated cell sorting (FACS) analysis of transduced cells. Consequently, we reduced the construct size by excising four of the 24 GCN4 peptide epitopes present in the original vector using two internal XhoI restriction sites. Additionally, we replaced the BFP fluorescence marker with an optimized puromycin resistance gene, facilitating efficient selection of transduced cells via puromycin treatment.

SLK cells were transduced with the newly modified SunTag-dCas9-20xGCN4 construct, pHRdSV40-NLS-dCas9-20xGCN4_v4-NLS-P2A-PuroR-dWPRE, and stable expression was selected using puromycin. Subsequently, we transduced SLK-SunTag-dCas9-20xGCN4 cells with lentiviral particles derived from pHR-scFv-GCN4-sfGFP-GB1-NLS-dWPRE, which encodes a GCN4-specific scFv antibody fragment fused to superfolder GFP (sfGFP)^8^. A low multiplicity of infection (MOI) was used to ensure minimal GFP expression, thereby optimizing the signal-to-noise ratio achievable with the SunTag system.

Next, we designed four guide RNAs (gRNAs) using standard gRNA design tools, targeting the 800 bp terminal repeat (TR) region of KSHV, which is present in approximately 40 copies per episome. Utilizing repetitive target elements is a critical requirement for dCas9-fluorescence-based imaging systems to substantially amplify target signals above background levels. Importantly, the terminal repeat units serve as latent origins of licensed DNA-replication and harbor discrete binding sites for the KSHV-encoded latency-associated nuclear antigen (LANA)^11–14^. Since LANA binding at these sites is of fundamental importance for episomal maintenance, the gRNA target sites were positioned at least 64 base pairs away from the previously defined binding sites (LBS1, LBS2, and LBS3)^11,12^.

Previous studies have demonstrated that the stem-loop sequence linking the tracr-RNA and gRNA into a single guide RNA (sgRNA) significantly influences the targeting efficiency of dCas9- and Cas9-based imaging and editing systems^15,16^. We adopted an extended stem-loop and A-U flip sgRNA sequence, which was reported by Chen and colleagues to enhance signal-to-noise ratios in dCas9-mCherry targeting of human telomeres^15^. Each TR gRNA sequence was initially cloned into a pCR2.1 construct containing a U6 promoter, BbsI-based gRNA cloning sites, and the optimized tracr-RNA sequence. The complete expression cassette was subsequently inserted into a LeGO-Zeo2 vector^17^ to generate lentiviral particles, thereby enabling stable sgRNA expression. An overview of the SunTag-KSHV system is provided in Figure 1A.

**Figure 1:**
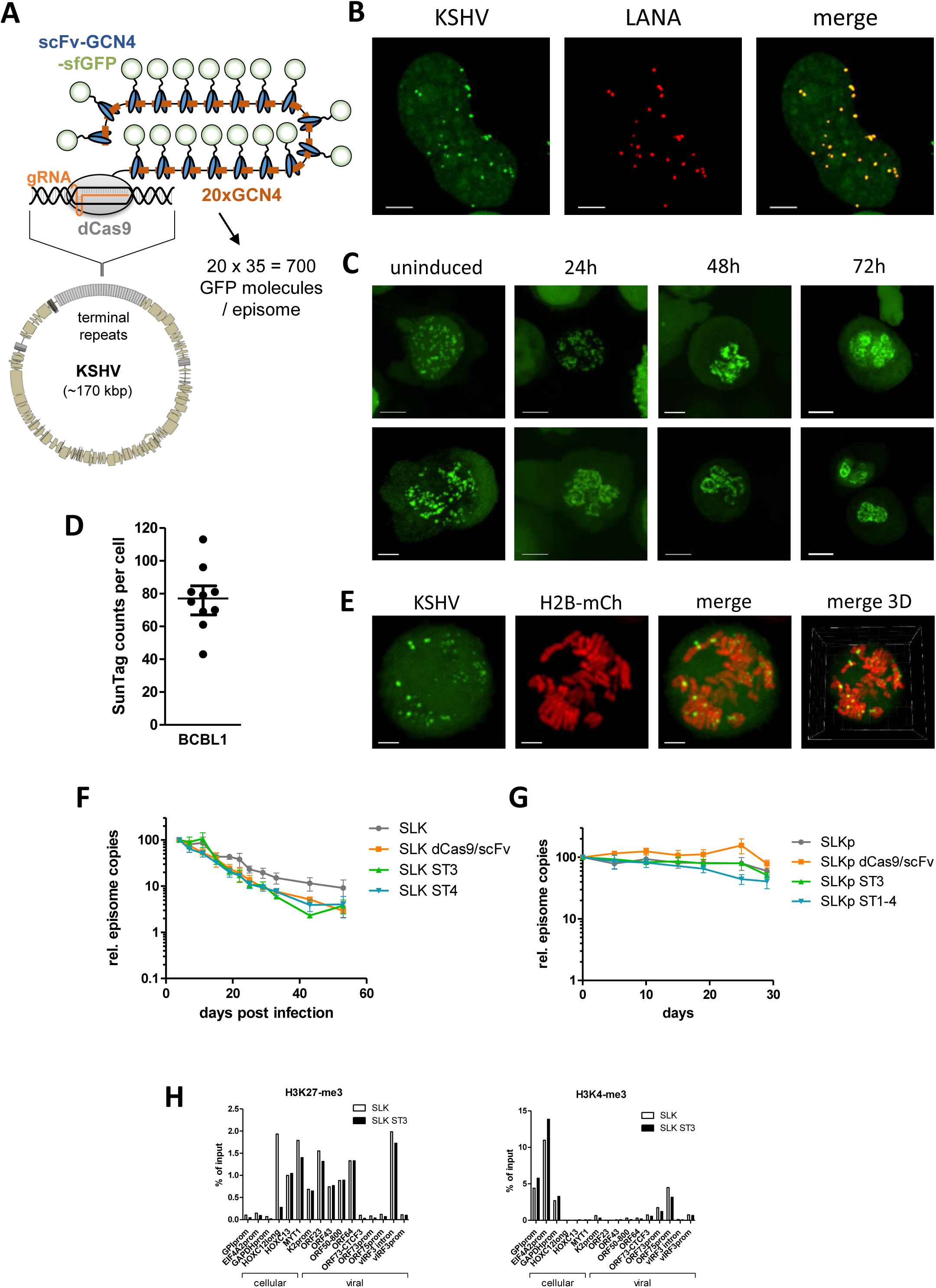
Establishment and characterization of a SunTag KSHV DNA live cell imaging system. A) Schematic model of the SunTag KSHV detection system. B) SLK-ST3 cells were infected with KSHV and fixed at 48 hours post infection with KSHV and fixed with paraformaldehyde to preserve the GFP fluorescence of the SunTag. LANA was stained by immuno-fluorescence. Images are represented as MIP of Z-stacks. Diffuse green background signal indicates the nucleus. Yellow dots in the merge image represent colocalizing signals. Scale bar = 3µm. C) SunTag KSHV visualization of lytic KSHV replikation in BCBL1 cells. Lytic replication of BCBL1-ST3 bulk cells was induced by treatment with TPA/Na+Butyrate. Images were acquired from uninduced cells and at the indicated time points post induction. Shown are two acquired images for each time point. Scale bar = 3 µm. D) A total number of 10 uninduced BCBL1-ST3 cells were used to measure the number of episomes per cell that are detectable by SunTag KSHV (n=10, median and interquartile range). E) LeGO-H2B-mCh transduced BCBL1-ST3 cell showing condensed chromosomes and KSHV episomes during mitosis. F) Quantification of KSHV episome loss in SunTag KSHV cells during latent infection. Relative KSHV episome copy numbers were normalized to the genomic locus of human GAPDHpro, three qPCR data sets of three biological replicates. Copy numbers of 4dpi were set to 100% for each cell line. Data are represented as mean +-SEM; SLK = plain SLK cells, SLK dCas9/scFv = SLK cells transduced with dCas9-GCN4 and scFv-GFP constructs, SLK ST3 = single cell clone SLK SunTag sgRNA TR3 clone 3, SLK ST4 = single cell clone SLK SunTag sgRNA TR3 clone 4. G) Relative KSHV episome copy numbers over time in SLK_P_ SunTag cells normalized to the genomic locus of human GAPDHpro (n=3). Data are presented relative to day 0 to indicate episome number changes over time and data are given as mean +-SEM. SLKp = latently KSHV infected cell line; SLKp dCas9/scFv = SLKp cells transduced with dCas9-GCN4 and scFv-GFP constructs; SLKp ST3 = bulk population of SLK SunTag cells with sgRNA TR3 construct; SLKp ST1-4 = bulk population of SLK SunTag cells transduced with mixed lentiviral particles of sgRNA TR1, 2, 3 and 4 expression constructs. H) ChIP-qPCR of H3K4-me3 and H3K27-me3 at cellular and viral target regions. GPIpro, EIF4A2pro and GAPDHpro serve as cellular positive controls for H3K4-me3 and as negative controls for H3K27-me3. HOXC12, HOXC13 and MYT1 serve as cellular positive controls for H3K27-me3 and negative controls for H3K4-me3.

SLK-dCas9-20xGCN4-scFv-GFP cells were transduced with the TR-specific sgRNA constructs and selected using zeocin. Bulk-transduced cell populations were designated SLK-ST1, SLK-ST2, SLK-ST3, and SLK-ST4 according to their respective sgRNA. These populations were infected with KSHV, and dot-like structures suggestive of KSHV DNA were monitored using conventional light microscopy. While all sgRNAs facilitated KSHV genome visualization, SLK-TR3 exhibited the highest signal accumulation in dot-like structures with an optimal signal-to-noise ratio (Figure S1). However, bulk populations displayed significant cell-to-cell variability in the presence of dot-like structures, likely due to differences in expression levels and stoichiometric ratios of scFv-GFP, dCas9-20xGCN4, and sgRNA-TR3.

Thus, we generated 46 single cell clones from the SLK-ST3 bulk population of sgRNA-TR3, infected them with KSHV and monitored the appearance of nuclear dot like structures upon infection. A total of eight clones were superior in their capability to produce dot like structures upon *de novo* infection as judged by conventional fluorescence light microscopy (Figure S1). The best detection capability was found in SLK-ST3 clones #3 and #4 (SLK-ST3#3 and SLK-ST3#4), thus, we used either a pool of the aforementioned eight clones or clones #3 and #4 individually for subsequent experiments as indicated.

To address this variability, we derived 46 single-cell clones from the SLK-ST3 bulk population expressing sgRNA-TR3. These clones were infected with KSHV, and nuclear dot-like structures were examined via fluorescence light microscopy. Eight clones demonstrated superior capability in generating dot-like structures upon *de novo* infection (Figure S1). The most effective detection was observed in clones SLK-ST3#3 and SLK-ST3#4, prompting us to utilize either a pooled selection of these eight clones or clones #3 and #4 individually for subsequent experiments, as specified.

### The SunTag KSHV System Enables the Study of All Phases of the KSHV Life Cycle

To confirm that the SunTag system specifically detects KSHV episomes, we first examined whether SunTag KSHV signals colocalize with LANA dots at 48 hours post-infection. It is well established that once sufficiently expressed, LANA binds to specific sites (LBS1-3) and accumulates at the terminal repeats of KSHV in dot-like structures, serving as a reliable marker of viral DNA^18^ at later stages of latent infection. We infected SLK-ST3 cells with KSHV and performed immunofluorescence (IF) analysis for LANA at 48 hours post-infection (Figure 1B, see Figure S2 for additional images and Movie S1 for 3D reconstruction). The SunTag-KSHV and LANA signals showed near-complete colocalization (yellow signal in the merged image).

After confirming that the SunTag KSHV system reliably detects and visualizes viral episomes, we explored whether it could also be used to observe viral replication centers after lytic. Since SLK cells exhibit limited support for lytic replication, we introduced the components of the SunTag-KSHV sgRNA#3 detection system into BCBL1 cells, which robustly support lytic replication upon induction. Similar to SLK-ST3 cells, BCBL1-ST3 cells exhibited nuclear dot-like structures (Figure 1C and Figure S2, and Movie S2). Using Imaris software, we quantified nuclear dots per individual BCBL1 cell, obtaining a median of 77 detectable dots per cell (Figure 1D), consistent with previously reported KSHV copy numbers per cell. Induction of the KSHV lytic cycle with TPA/Na+ butyrate resulted in the formation of tube-like structures, resembling the characteristic herpesviral replication compartments (Figure 1D, Figure S2, Movie S3, Movie S4 and Movie S5).

Although the primary aim of developing an episome tracking and visualization system was to monitor the early stages of KSHV infection—when traditional detection techniques like LANA IF are not applicable—we also investigated its potential for long-term latency studies. The SunTag gRNAs were designed at least 64 bp up- or downstream of the nearest LBS to minimize interference with LANA binding; however, the relatively large SunTag complex could still potentially impact LANA’s tethering function. To assess whether SunTag KSHV affects episome tethering, we transduced BCBL1-ST3 cells with a lentiviral H2B-mCh expression vector and imaged cells with condensed mitotic chromosomes (Figure 1E and Movie S6). All detectable SunTag KSHV signals localized near or in direct contact with mitotic chromosomes, as previously demonstrated via LANA immunofluorescence in mitotic KSHV-infected cells, indicating that SunTag KSHV does not impair LANA-mediated tethering. However, prolonged tracking of infecting episomes might slightly reduce tethering efficiency.

To determine whether SunTag KSHV impacts latency establishment and episome maintenance, we infected two SLK-ST3 clones (#3 and #4), the parental SLK-dCas9-GCN4 cell line, and untransduced SLK cells with KSHV in triplicate experiments and monitored episome content over 56 days via qPCR (**Figure *1***F). While we observed no significant differences in episome loss or final copy numbers between the two SunTag clones and the parental dCas9 cell line, a slight reduction in copy numbers was detected compared to SLK cells, suggesting a minor impact from the dCas9 or scFv-sfGFP construct.

In addition to *de novo* infection studies, we investigated whether introducing the SunTag KSHV system into cells with pre-established latency affects episome maintenance and tethering. Since BCBL1 cells rely on latent KSHV for survival, episome loss would generate selection pressure, introducing bias into the analysis. To circumvent this, we introduced the SunTag KSHV system (sgRNA#3 and sgRNA#4) into a previously described KSHV-maintaining SLK cell population called SLK ^19^, which does not require KSHV for viability. Similar to the earlier experiment, we compared episome content in SLK_P_-ST3 and SLK_P_-ST4 cells with their parental counterparts via qPCR over 30 days in triplicate experiments (Figure 1G). Consistent with our earlier findings, we detected no differences in copy numbers.

To rule out potential interference of SunTag KSHV with chromatinization processes that could alter epigenetic modifications, we performed ChIP-qPCR for H3K4-me3 and H3K27-me3 at two days post-infection (Figure 1H). The presence of the SunTag KSHV system did not affect the levels or patterns of these histone modifications in repressed or active regions of the episome.

Considering all findings, we conclude that the SunTag KSHV system enables reliable detection of KSHV episomes across all phases of the viral life cycle, facilitating the study of *de novo* infection, mitosis, latency, and lytic replication.

### Development of the <u>Sun</u>Tag <u>S</u>ingle-V<u>e</u>ctor <u>T</u>argeting System (SunSeT)

As previously mentioned, generating cell lines or clones through consecutive transduction with multiple SunTag components limits the application of the original system to tumor or immortalized cell lines, as these allow multiple rounds of transduction and selection with two antibiotics and GFP sorting. Moreover, controlling the expression ratios of individual components is challenging, and multiple transduction steps may interfere with the host innate immune response. To overcome these limitations, we utilized the piggyBac vector system, which enables transposase-mediated integration of target sequences into the host genome via simple transfection or electroporation of the piggyBac construct alongside a transposase expression vector^9,10^. The transient expression of transposase ensures stable integration of the long terminal repeat (LTR)-flanked target sequence within the piggyBac vector.

We combined all SunTag system components into a single piggyBac vector, as illustrated in Figure 2A. The high cargo capacity of piggyBac allowed the inclusion of three separate expression cassettes for sgRNA-TR2, sgRNA-TR3, and sgRNA-TR4, along with the 24xGCN4 peptide array fused to P2A-PUROopt, enabling puromycin selection of integrated constructs. Additionally, we replaced the original promoter with an intron-containing CAG promoter to enhance mRNA stability and mitigate cellular responses to intronless long RNAs.

**Figure 2:**
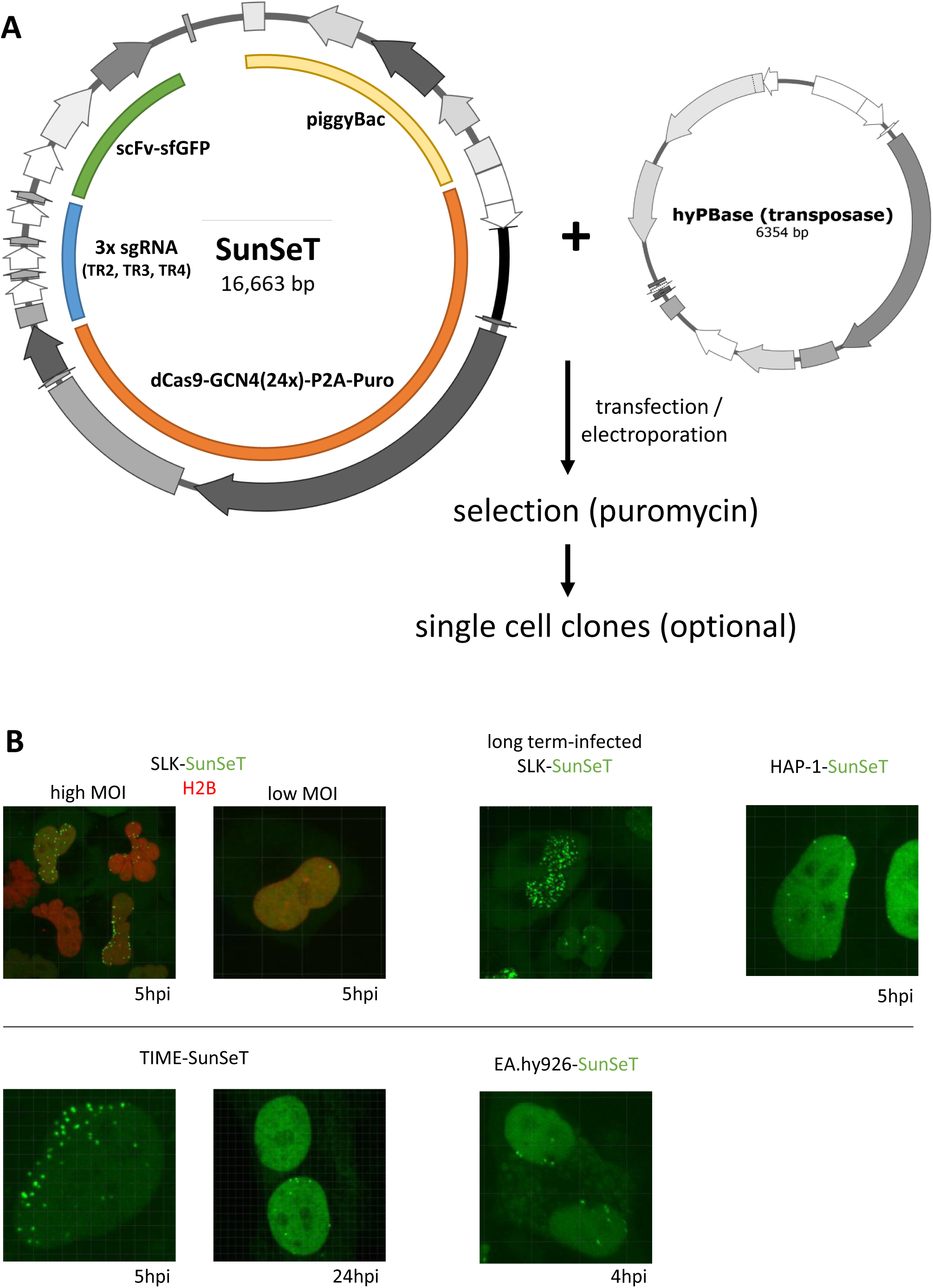
Generation of a SunTag Single-Vector Targeting System (SunSeT) A) Schematic view of the SunSeT vector and transfection system containing all components of the original SunTag system as well as three sgRNA expression cassettes for sgRNA-TR2, -TR3 and TR-4. B) Different cell lines were electroporated with SunSeT and transposase vectors and selected by puromycin 48 hours after electroporation. SLK were additionally transduced with an H2B-mCherry construct, to visualize chromatin in the nucleus. Cell lines were infected with KSHV and imaged at the indicated time points. Long term KSHV infected SLK cells were imaged after 5 days of puromycin selection.

We successfully introduced the SunSeT system into a diverse range of cell types, including tumor cell lines (SLK, HAP-1), long-term KSHV-infected SLK cells, the hybrid EA.hy926 cell line (derived from the fusion of HUVEC and A549 cells)^20^, and telomerase-immortalized microvascular endothelial cells (TIME) (Figure 2B). Both EA.hy926 and TIME are widely used model cell lines in KSHV research.

### SunTag-KSHV Reveals Limited Early Nuclear Localization Dynamics During *de novo* Infection

After validating that the SunTag KSHV system effectively visualizes KSHV episomes in living cells, we examined the early stages of *de novo* infection. This critical phase of the viral life cycle involves key processes such as episomal circularization, deposition of core histone proteins by histone chaperones, chromatinization, and binding of cellular transcription factors and repressor complexes, which collectively shape and stabilize the latent expression profile.

We infected SLK-ST3 cells with KSHV and immediately monitored the spatiotemporal distribution of SunTag KSHV signals in the nucleus using high-temporal-resolution imaging, acquiring Z-stack images every five minutes (Figure 3A, Movie S7 and Movie S8). The first detectable KSHV signals appeared near the nuclear membrane within 20 to 30 minutes post-infection. The number of detectable signals increased over time, stabilizing approximately two hours post-infection, with only a slight increase in individual dot intensity. This suggests that most viral DNA is released from capsids into the nucleus within this timeframe. However, given that SunTag recruitment requires binding to viral DNA, we estimate a detection delay of up to 20 minutes before signals rise above background levels.

**Figure 3:**
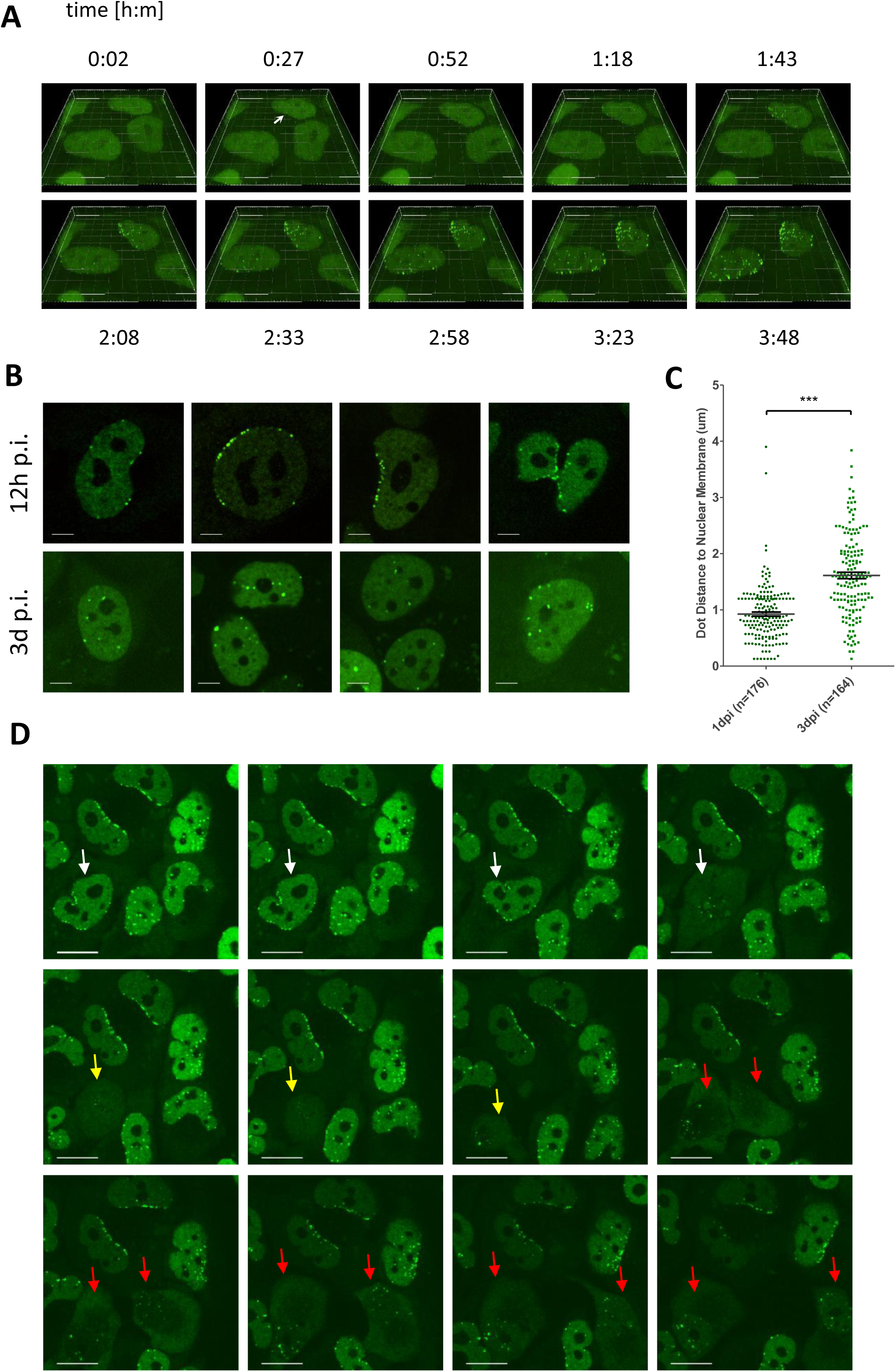
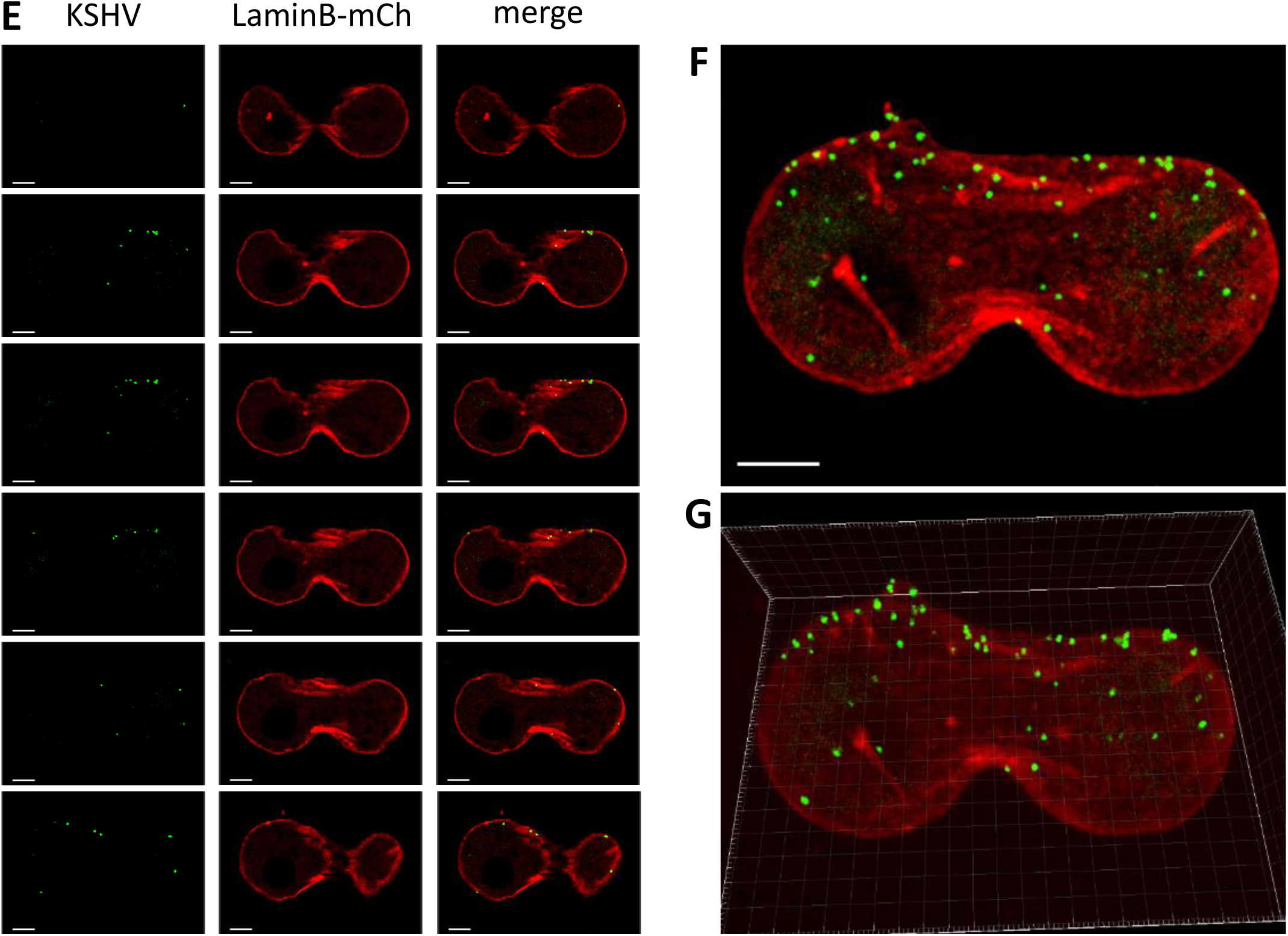
Early episome dynamics and localization detected by SunTag. A) SLK-ST3 cells were infected with KSHV and monitored by 3D live cell imaging during the first 4 hours of infection. Images represent 3D reconstructions of Z-stacks at the indicated time points. The arrow indicates the first detectable signal at ∼30 min post infection. Images represent a subset of a time series (see movie S07) with image acquisition every 5 minutes over 4 hours, thus, indicating a high stability of the SunTag KSHV signal throughout repetitive sample exposures. B) Confocal images of SLK-ST3 cells at 12h (upper panel) and 3d post infection (lower panel) indicating the KSHV episome localization in pre and post mitotic cells. Scale bar = 3µm. C) Distance of SunTag KSHV dots to the nuclear membrane as determined by Imaris. Green nuclear sfGFP fluorescence was used to define the nuclei at 24h and 3d p.i.. (mean +-SEM). D) SunTag KSHV signals throughout mitosis. Shown is a time laps series of mitosis (left to right and top to bottom). Arrows indicate the pre-mitotic (white arrows), mitotic (yellow arrows) and the two post mitotic daughter cells (red arrows). Scale bars = 10 µm (see movie S09). E-G) LaminB-mCh expressing cells were imaged 8 hours post infection with KSHV. Localization of KSHV episomes at the nuclear lamina is detectable in confocal Z-stack images (E) as well as in MIP (F) and 3D reconstruction (G).

Interestingly, viral DNA entry predominantly occurred on one side of the nucleus. Given the kidney-like shape of many SLK nuclei, we hypothesize that entry occurs preferentially near the microtubule organizing center (MTOC), consistent with previous reports on microtubule-mediated viral transport to the nuclear pores^21,22^.

Notably, KSHV episomes exhibited minimal movement from their entry site over the four-hour imaging period, suggesting an absence of active intra-nuclear transport mechanisms. However, after several days of infection, we observed an increasing number of cells with episomes distributed further from their initial nuclear entry sites (Figure 3A, B and D). Distance measurements of individual signals relative to the nuclear membrane (defined by nuclear GFP signal boundaries) one and three days post-infection revealed a significant increase in median distance (Figure 3C). As no active episomal movement was detected, we hypothesize that cell division plays a key role in redistributing episomes throughout the nucleus.

We further tested this hypothesis through live-cell 3D imaging of mitotic cells (Figure 3D, Movie S9). Pre-mitotic cells harbored episomes predominantly at the nuclear periphery. During mitosis, episomes dispersed throughout the nucleus. Since chromosome territories remain stable during mitosis, this suggests that LANA-mediated episome tethering does not permanently fix episomes to specific nuclear sites. Instead, tethering appears to be dynamic, with episomes gaining mobility during chromosome condensation and subsequently attaching to mitotic chromatin fibers.

Although the majority of episomes in our initial experiments remained localized near the lamina until the first cell division, a minor fraction was observed at a virtually more distal location immediately after infection. While it appeared formally possible that these episomes had migrated prior to dCas9 recruitment and SunTag detection, we considered it more likely these events result from episomes having entered the nucleus via invaginations and tunnels extending into the nucleoplasm.

To investigate whether episomes associate with nuclear membrane invaginations, we therefore expressed a LaminB-mCherry fusion protein in SLK-ST3 cells. Live-cell imaging confirmed that all *de novo* KSHV episomes colocalized with LaminB, confirming that episomes are retained at their nuclear entry sites after infection, thus, providing additional insights into early nuclear localization dynamics (Figure 3E-G, Movie S10 and Movie S11).

### Long-Term Tracking of KSHV Episomes Through Multiple Cell Divisions

Having established robust visualization of KSHV episomes in both *de novo* infected and stably infected cells, we next assessed the system’s performance during long-term imaging across multiple days and cell divisions. To achieve this, we utilized SunSeT SLK cells, which stably harbor high-copy numbers of latent KSHV episomes. To facilitate cell tracking, these cells were further transduced with H2B-mCherry, allowing for precise visualization of individual cells. The heterogeneous expression levels of H2B-mCherry also enhanced tracking of individual cells over extended periods.

We conducted live-cell imaging over a five-day period and successfully monitored a single cell through three successive divisions (Figure 4A and Movie S12). Automated quantification of episome copy numbers in the parental and daughter cells at multiple time points demonstrated stable episome retention up to the third generation (Figure 4B and C). However, due to limitations in the imaging field and the occasional loss of cells from the focal plane, episome quantification beyond the third generation was not possible for all cells.

**Figure 4:**
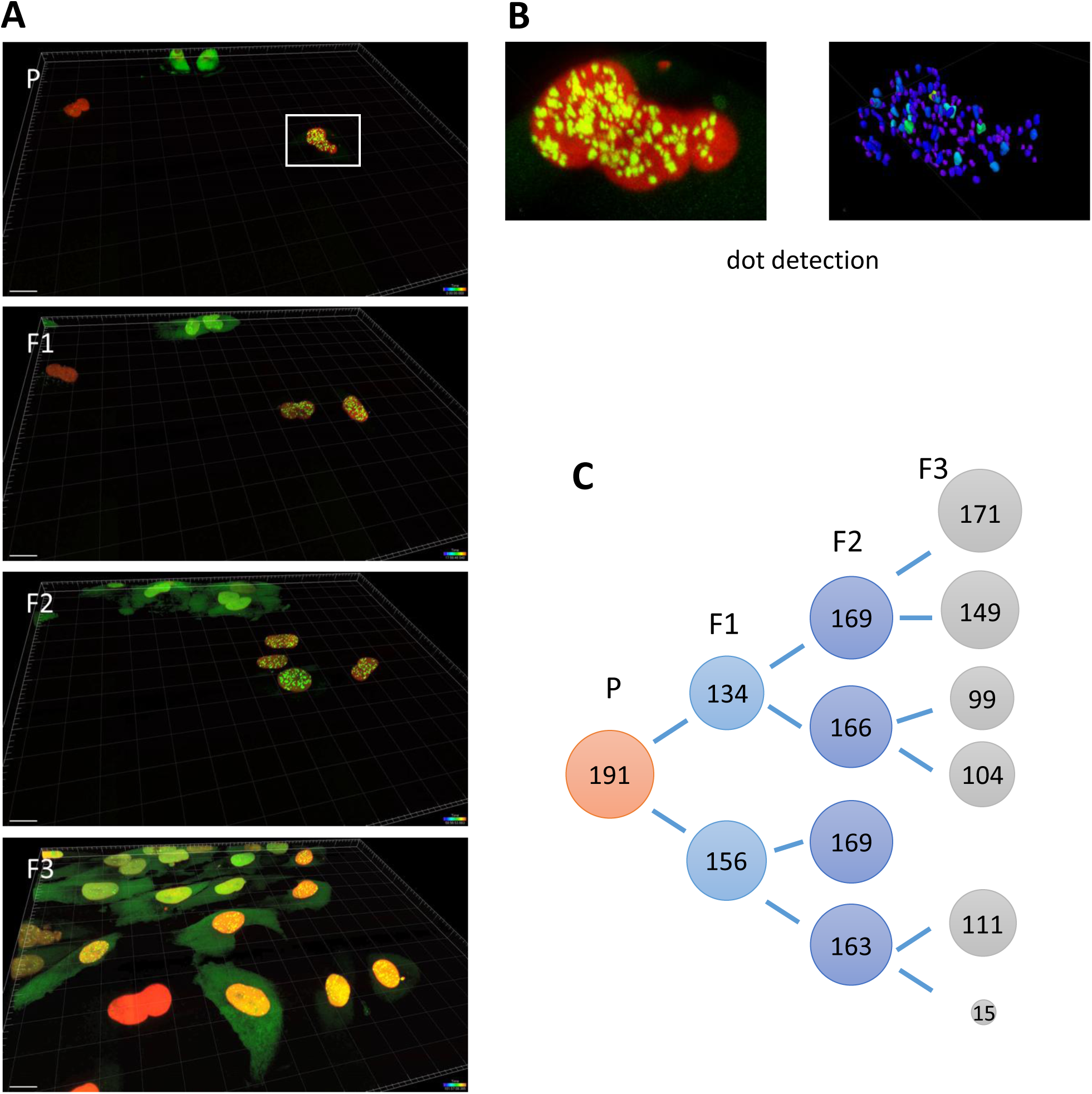
Long-term SunTag KSHV imaging over several rounds of cell division. A) Long-term KSHV infected SunSeT-SLK cells were transduced with H2B-mCherry to visualize host chromatin. Z-stack images of cells were taken with a spinning disk microscope every 2 hours over a total period of 5 days. Shown are selected images from four cell generations (P to F3). B) Individual dots were detected and quantified using Arivis analysis Software. C) Arivis quantification of SunSeT signals along cell division trajectories was performed for all images. Shown are the maximum counts of all images taken from the respective cells.

### KSHV Does Not Localize to PML-NB Structures During the Early Stages of Infection

We demonstrated previously that KSHV does not appreciably associate with promyelocytic leukemia protein (PML) nuclear bodies (PML-NB, also referred to as Nuclear domain 10 or ND10 bodies) during early or late stages of latent infection^23^. Our earlier analyses, however, were based on immunofluorescence experiments evaluating co-localization between LANA and PML or SP100, key components of PML-NB. In cells with established latency, robust binding of LANA to viral terminal repeats has been shown to result in formation of nuclear LANA dots that represent a valid proxy for single episomes or episome clusters^24^. However, as LANA is not a component of virions it needs to be newly expressed from incoming episomes, causing a significant delay before LANA dots become apparent (approximately 4-8 hours until first dots appear, with the total number of dots continuously increasing until 24 to 48 hours).

Considering the above, we considered it possible that viral episomes may transiently interact with PML-NB prior to LANA dot formation. Likewise, it appeared formally possible that PML-NB association may prevent LANA-dot formation on a subfraction of episomes, potentially leading to subsequent episome sequestration or transcriptional silencing. Using the SunTag KSHV system, we therefore examined potential colocalization during earlier stages of infection via live-cell imaging, using either an SP100-mCherry or a PML-IV-mCherry fusion construct to mark PML-NB.

Consistent with our previous findings, we did not observe colocalization of KSHV episomes with PML-NB at any time point - neither during the early phase of infection, nor at later stages when LANA was already expressed (Figure 5, Movie S13 and Movie S14). While our data thus confirm and extend our previous findings, it should be noted that experiments were performed with a fluorescently marked PML-IV isoform. We expect this isoform to accurately mark canonical PML-NB, but it remains possible that specialized PML-IV-negative PML-NB, or isoforms not residing in PML-NB could interact with *de novo* infecting KSHV episomes. We are currently performing additional KSHV-SunTag experiments to evaluate these possibilities.

**Figure 5:**
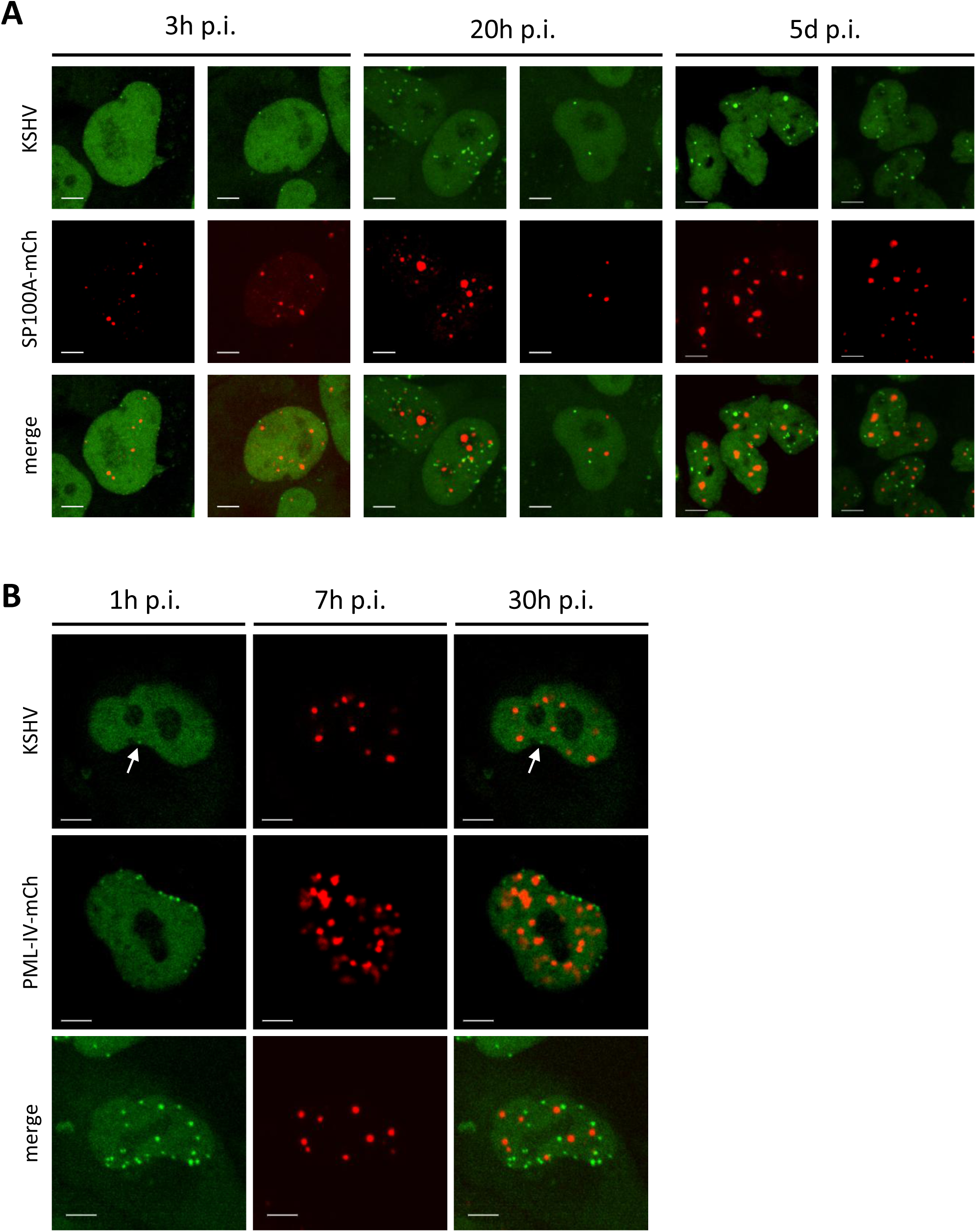
*De novo* infecting KSHV episomes do not colocalize with PML-NBs. SLK-ST3 #3 were transduced with either a lentiviral SP100A-mCh (A) or PML-IV-mCh (B) expression vector and after sorting for mCherry expression were infected with KSHV. Images were acquired during live cell imaging at the indicated time points. Shown are MIP images (scale bar = 5µm).

### SunTag KSHV Reveals Transient IFI16 Interactions with Individual Episomes

IFI16 is a well-established DNA sensor that recognizes herpesviral DNA upon infection and transfection ^25–28^. However, direct visualization of IFI16 interactions with viral DNA has been limited to fixed-cell analyses, such as fluorescence in situ hybridization (FISH), or indirect detection using viral DNA-associated proteins like LANA. These methods inherently lack temporal resolution, preventing a dynamic view of IFI16–viral DNA interactions.

Using our SLK SunTag KSHV system combined with IFI16-mCherry expression in live-cell imaging, we visualized the temporal dynamics of IFI16-mediated DNA sensing during *de novo* KSHV infection. In uninfected SLK cells, IFI16 is predominantly and uniformly localized within the nucleus.

Consistent with previous studies, we observed a multiphasic redistribution of IFI16 following KSHV infection^25^. Early in infection (as soon as 1 hour post-infection), IFI16 signals accumulated at the nuclear lamina, colocalizing with SunTag KSHV signals (Figure 6A). Notably, IFI16 accumulation at the perinuclear region preceded the detection of SunTag signals (Figure 6B). During this early phase, individual IFI16 puncta associated with single SunTag KSHV signals, indicating efficient sensing of individual herpesviral episomes by the host (Figure 6B).

**Figure 6:**
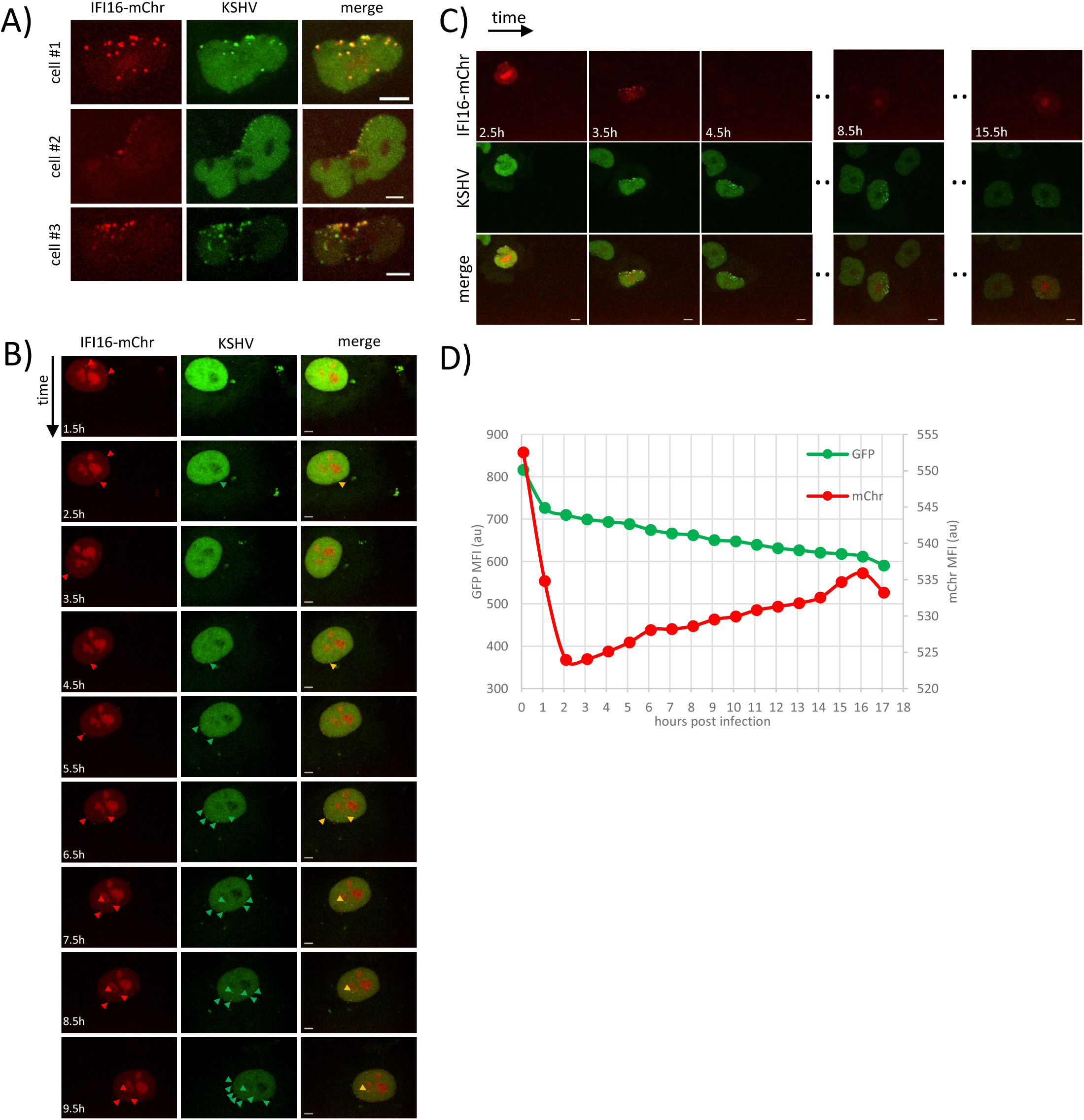
SunTag KSHV visualizes transient interaction of IFI16 with *de novo* infecting episomes. SLK-ST3 #3 cells transduced with a lentiviral IFI16-mChr expression construct were infected with KSHV and subjected to time lapse live cell imaging. A) Representive cells showing co-localization of IFI16-mChr with SunTag KSHV signals. MIP images, scale bar 5 µm. B) Time lapse imaging during KSHV infection reveals early and transient perinuclear IFI16 signals. Arrow heads indicate IFI16 (red), KSHV (green) and co-localized (yellow) signals. MIP images at the indicated time points (scale bar = 5µm). C) Time lapse imaging during KSHV infection reveals IFI16-mChr degradation after appearance of perinuclear puncta. MIP images at the indicated time points (scale bar = 5 µm), MFI Mean Fluorescence Intensity.

Following initial DNA sensing, IFI16 puncta rapidly disassembled, while SunTag KSHV signals remained visible. This process appeared synchronized at the level of individual infectious particles, suggesting that IFI16 engagement with each KSHV episome is transient and resolved individually, likely through host or viral regulatory mechanisms. Furthermore, in cells infected with a high number of KSHV particles, IFI16 degradation was evident, as indicated by a sharp reduction in mCherry fluorescence at 2 hours post-infection (Figure 6C and D). This reduction was not due to fluorophore photobleaching, as GFP fluorescence remained stable throughout imaging, and mCherry fluorescence levels recovered after 2 hours due to continuous IFI16-mCherry expression.

Taken together, these results confirm previous findings on the rapid sensing of herpesviral DNA by IFI16 and its subsequent counteraction through IFI16 redistribution and degradation^25,28,29^. The consistency of these observations with prior studies underscores that the SunTag system does not interfere with the viral life cycle. Moreover, the SunTag KSHV system allows direct correlation of IFI16 perinuclear puncta with herpesviral DNA entry sites, exemplifying its utility for studying rapid protein-DNA interactions at the level of individual episomes.

### KDM2B Localizes to Viral Episomes Immediately After Nuclear Entry

Upon nuclear entry, the epigenetically naïve and non-chromatinized viral DNA is exposed to various nuclear factors. Our recent findings have demonstrated that the lysine-specific histone demethylase KDM2B binds to KSHV episomes early in infection^30^. KDM2B is primarily known for its role in binding CpG islands and preventing di-methylation of H3K36 at active promoters. However, more recently it has also been recognized as a modular PRC1 component that guides vPRC1.1 to CpG islands. Since our earlier analysis had suggested that KSHV exposes CpG island characteristics to promote rapid acquisition of facultative heterochromatin^30^, we sought to investigate temporal dynamics of KDM2B association with viral episomes. Using a KDM2B-mCherry fusion construct, we therefore monitored co-localization with KSHV-SunTag (Figure 7A, Movie S15, Movie S16 and Movie S17). While KDM2B was distributed throughout the nucleus (including the previously described nucleolar staining^31^, we observed increased accumulation near the nuclear membrane approximately 30 minutes post-infection, with KSHV episomes becoming detectable at the same sites simultaneously or shortly after KDM2B accumulation. Within the first two hours of infection, all episomes visibly colocalized with KDM2B, confirming its early interaction with KSHV (Figure 7F). This association remained stable at later stages of infection (Movie S18).

**Figure 7:**
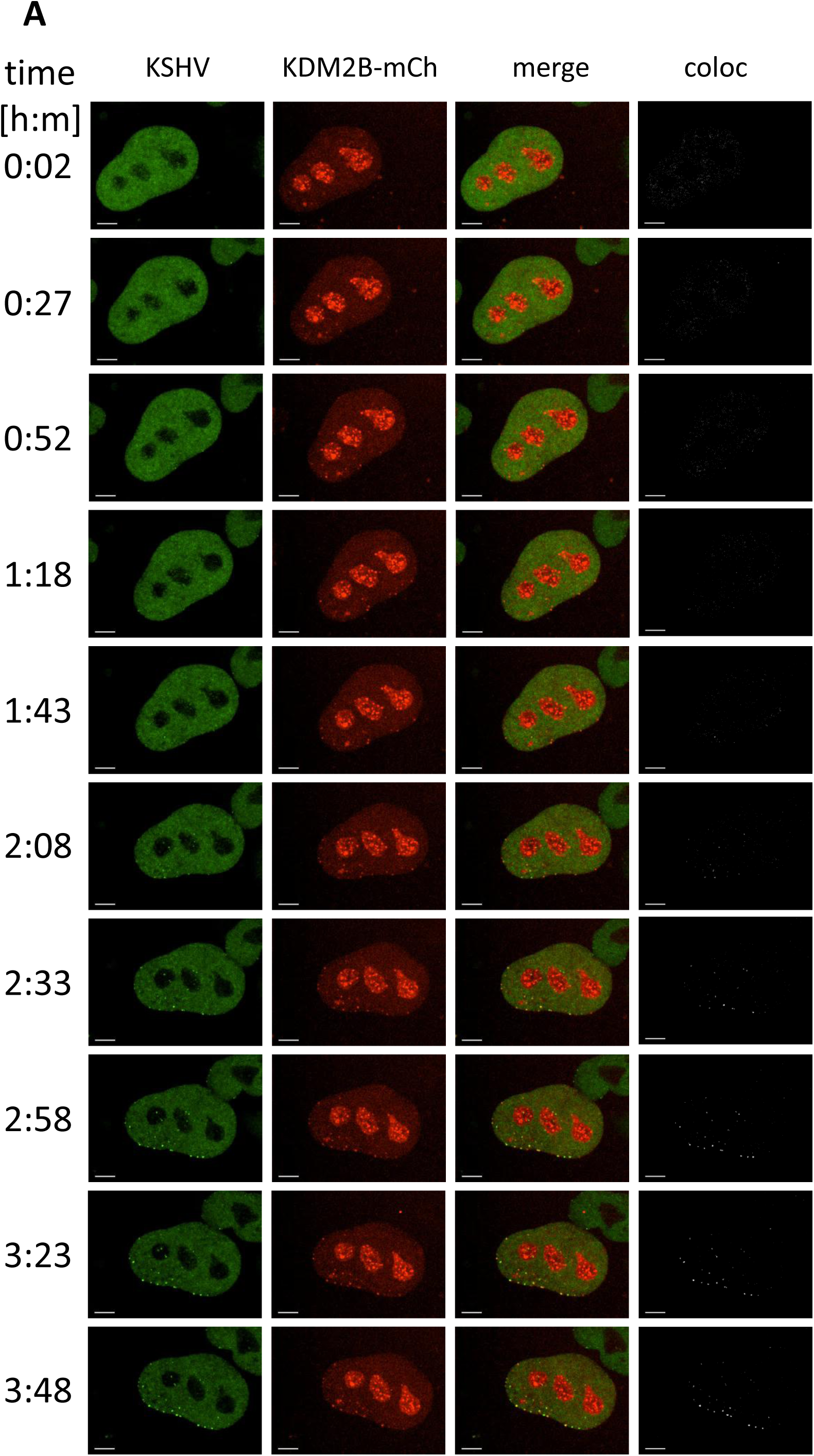
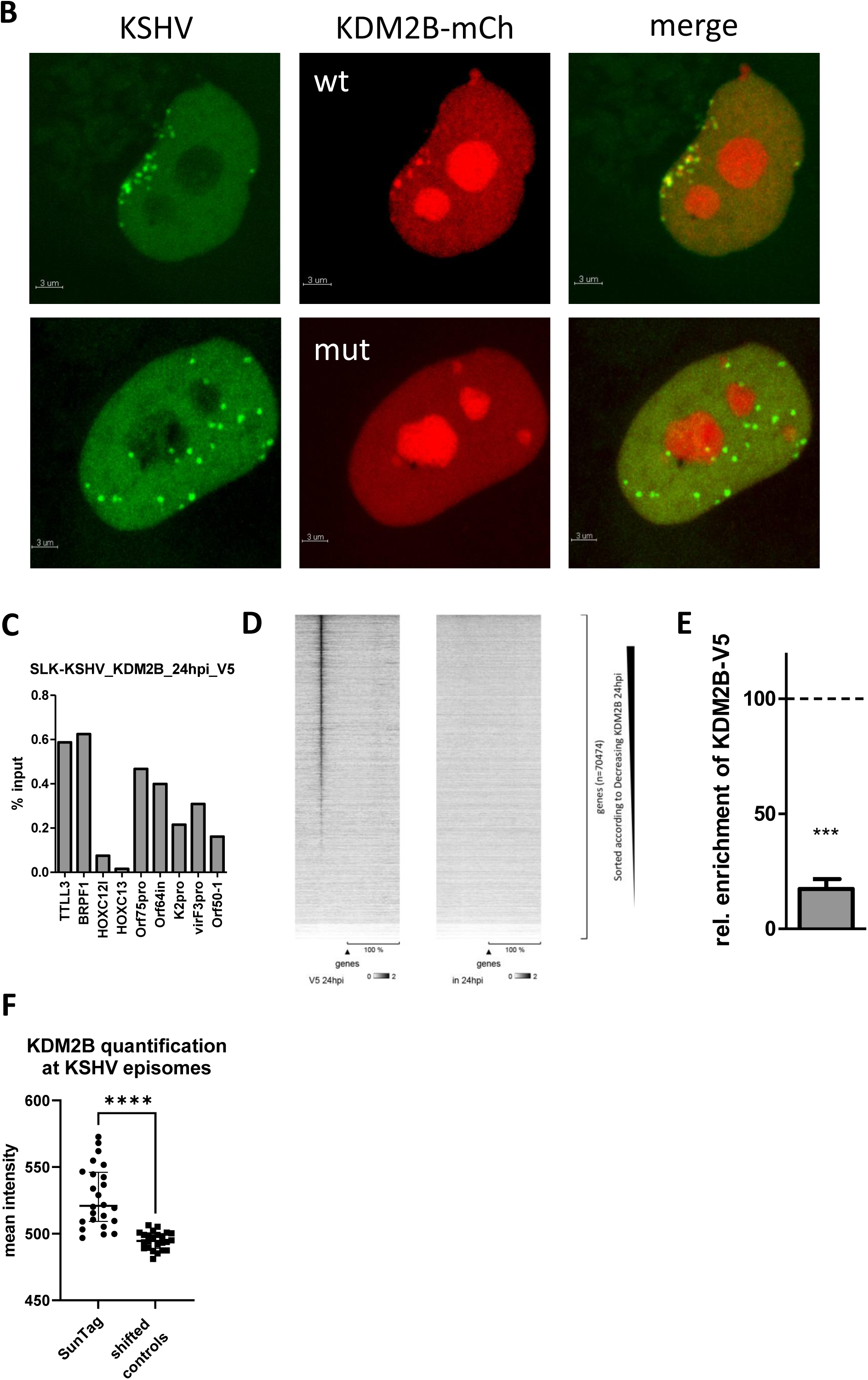
SunTag KSHV visualizes early binding of KDM2B to *de novo* infecting episomes. A) SLK-ST3 cells were transduced with a KDM2B mCherry fusion construct and infected with KSHV. Images were taken as a time laps movie every 5 minutes and are shown as MIP at the indicated time points post KSHV infection (scale bar = 3µm). B) Images of SLK-ST3 KDM2B-mCh and KDM2BmutZF-mCh were acquired 24h p.i. with KSHV and are shown as MIP of confocal Z-stacks. C) ChIP-qPCR analysis of KDM2B-mCh-V5 binding to cellular target loci and KSHV at 24 h post infection. D) ChIP-seq heatmap of KDM2B-mCh-V5 binding to human promoter regions at 25 hours post infection with KSHV. Input is shown as a control sample. E) KDM2B-mCh-V5 is significantly enriched on KSHV in comparison to host negative and positive control regions. Data is presented relative to host control regions. 100% corresponds to the median of the top 200 KDM2B-bound regions (according to padjust value) and 0% to the median of 200 negative sites (defined as ChIP background). F) Quantification of KDM2B-mCherry signal intensity at SunTag dots and at shifted background controls.

KDM2B contains a CXXC zinc finger domain that specifically recognizes and binds unmethylated cytosines within CpG contexts—an essential feature of its function. To determine whether KDM2B recruitment to episomes in the SunTag system was due to direct DNA binding rather than indirect interactions or mCherry aggregation, we generated a CXXC mutant in the KDM2B-mCherry fusion protein and assessed its recruitment capabilities. As shown in Figure 7B, the mutant failed to colocalize with KSHV episomes, strongly supporting the hypothesis that KDM2B binds to the high-density unmethylated CpGs present in KSHV.

To further validate that the mCherry fusion construct directly binds viral DNA, we introduced a double V5 tag into the KDM2B-mCherry construct and performed ChIP-qPCR and ChIP-seq at 24 hours post-infection in SLK cells (Figure 7C–E). ChIP-qPCR analysis confirmed strong binding of KDM2B-mCherry-2xV5 to cellular control loci TTLL3 and BRPF1, as well as all tested KSHV loci, with no detectable binding to negative control loci HOXC12 and HOXC13. Expanding the analysis to all human promoters revealed strong promoter association, consistent with previous studies^30,32^. Since no distinct binding pattern was observed early in infection, we calculated the overall enrichment of KDM2B-mCherry-2xV5 on KSHV episomes, using the 200 most significant cellular target loci and 200 background controls as described previously^30^. These analyses demonstrated that KDM2B-mCherry-2xV5 is significantly enriched on KSHV episomes compared to background levels.

## Discussion

Here, we describe the development and application of a CRISPR/Cas9-based 3D live cell imaging system capable of tracking single KSHV episomes in real-time which exploits the SunTag^8^ technology. We designed sgRNAs to target SunTag complexes to the terminal repeat units of KSHV, which resulted in high level signal amplification upon infection. Furthermore, we combined the original SunTag components into a piggyBac system (SunSeT) to facilitate rapid generation of a broad range of SunTag/SunSeT KSHV reporter cell lines.

Incoming viral genomes were detectable within the first hours of *de novo* infection and could be tracked via live cell imaging through different stages of the viral lifecycle including latency and lytic replication. We were able to investigate early and late localization of individual episomes as well as transient and stable association with host factors and demonstrated, that the SunTag itself does not interfere with the different stages of the viral life-cycle.

The SunTag system revealed that incoming viral DNA is retained at the nuclear lamina without indication of immediate nuclear transport or diffusion. This indicates that crucial early events such as circularization and early chromatinization occur within this subnuclear compartment. Whether or not unique properties of the lamina-proximal environment (e.g., enrichment of transcriptionally silent B chromatin compartments^33^ may promote establishment of latency remains to be investigated. Since we observed dispersion of KSHV genomes after mitosis, we expect that our system will enable studies of dynamic association with other subnuclear compartments or regions during latency. For example, Epstein-Barr virus (EBV) episomes have been shown to associate with AT-rich, gene-poor chromatin regions during latency^34^. Investigating whether KSHV follows a similar pattern could provide valuable insights into conserved herpesvirus latency mechanisms.

Using this system, we confirmed the previously reported absence of interaction between PML-NB and *de novo* infecting KSHV episomes^30^. Additionally, we demonstrated the ability to observe both transient and stable interactions between KSHV episomes and key cellular regulators, specifically IFI16 and KDM2B. Both interactions have been reported previously but became now quantifiable in a temporal dimension further substantiating their mode of interaction with KSHV genomes. In agreement with our previous ChIP-seq analysis^30^, SunTag data demonstrate that not only a subset, but indeed all episomes colocalize with KDM2B very early in infection, indicating a robust and universal association with KSHV which depends upon KDM2B’s ability to bind unmethylated CpG dinucleotides. This broad interaction underscores the previously proposed role of CpG island characteristics in KSHV genomes in PRC recruitment. In a more recent study, we furthermore investigated chromatin states adopted by latent KSHV episomes in cells with individual and combined knockouts of essential PRC1 and PRC2 components^35^. Our results confirm that KSHV represents a formidable PRC target, with viral genomes acquiring facultative heterochromatin marks primarily via the PRC1 axis. However, the fact that functional knockout of PRC1 resulted in reduced, but not abolished acquisition of the PRC2-dependent histone mark H3K27me3 indicates that PRC recruitment can also proceed via other routes, most likely the PRC2.1 component PCL1-6^36^ or other CpG-island binding components of modular PRCs (reviewed in ^37^).

Interestingly, our PRC knockout study also demonstrated that, in the absence of PRCs, KSHV episomes become subject to transcriptional silencing by the human silencing hub (HUSH) complex^35^. While these results suggest a protective role of PRCs against HUSH, a recent study by Roubille and colleagues reported HUSH-mediated H3K9me3 deposition on PML-NB-associated HSV-1 genomes. Our observation that incoming KSHV episomes fail to associate with PML-NB suggests that KSHV may evade these structures to adopt favorable facultative heterochromatin states. Considering that KSHV appears to differ substantially from HSV1^38^ and other DNA viruses that readily associate with PML during *de novo* infection (reviewed in ^39,40^), it will be of fundamental interest to identify underlying recruitment modalities. We expect that single-molecule imaging systems such as the one described here will be of considerable value in elucidating such mechanisms.

Inflammasome induction and IFI16 colocalization with KSHV has been reported previously^27^ and IFI16 degradation is an established observation in HSV-1 infection. More controversial are the factors that target IFI16 for proteasomal degradation. There is evidence for HSV-1 ICP0 mediating IFI16 degradation, but this has been challenged^28,29^. The factor(s) for KSHV-mediated IFI16 degradation are still elusive, although a late lytic gene product has been suggested as IFI16 is also degraded upon re-activation from latency^41^.

The adaptability of the SunTag/SunSeT-KSHV system makes it a broadly applicable tool for studying not only KSHV but also other targets containing repetitive elements. By allowing the cloning of custom sgRNAs into the backbone, this system can potentially be tailored to investigate e.g. other viruses or host factors involved in episome regulation. Recent advances in dCas9-based imaging already allow the detection of single loci or even multiple targets by multiplexing of single locus guide RNAs like the CRISPR/Casilio-based imaging method^42^. Adaptation of similar strategies into the SunSeT format might further improve detectability of viral DNA.

Taken together, our findings demonstrate that the SunTag-KSHV system is a powerful tool for investigating the spatial and temporal dynamics of viral genomes in living cells. The ability to track episome behavior over time provides novel insights into herpesvirus nuclear interactions, host factor engagement, and potential mechanisms governing viral latency and persistence. The system’s capacity to visualize viral-host interactions in real-time offers a significant advancement in the study of herpesvirus biology and approaches seeking to develop new avenues for targeted intervention.

## Materials and Methods

### Cell lines

BCBL1 cells ^43^ were cultured in RPMI 1640 medium containing 10% FCS, 100 U/ml Penicillin and 100 μg/ml Streptomycin. SLK cells have been recently discovered to be a misidentified cell line (Cellosaurus AC: CVCL_9569) and was identified as a contaminating cell line (Caki-1; Cellosaurus AC: CVCL_0234) and is listed in *ICLAC.* Nevertheless, these cells have been shown to efficiently support KSHV latency as stated in several publications throughout the last years and therefore represent a suitable model cell line to study chromatinization of latent KSHV genomes. SLK_P_ cells represent a previously described pool of stably long-term infected single cell clones derived from SLK cells infected with the BCBL1 KSHV strain ^19^. All SLK derivates were grown in DMEM High Glucose (Gibco, Darmstadt, Germany) supplemented with 10% FCS, 2 mM L-Glutamine, 100 U/ml Penicillin and 100 μg/ml Streptomycin. After transduction with PuroR or ZeoR containing lentiviral expression constructs, culture media were supplemented with 0.5µg/ml puromycin (Invitrogen) and/or 200µg/ml Zeocin (Invitrogen), respectively. EA.hy926 ^20^ (ATCC: CRL-2922) and TIME cells (ATCC: CRL-4025) were cultured according to the vendors recommendations.

### KSHV virus stock preparation and infection

Infectious wild-type KSHV supernatant was produced by inducing latently infected BCBL1 cells with TPA (Sigma-Aldrich; Cat#P8139-5MG) and sodium butyrate (Sigma-Aldrich; Cat#B5887-1G) followed by 100x concentration by centrifugation as described previously ^199^. For infection, concentrated KSHV stock solutions were diluted in 600 µl EBM-2 medium without supplements (LONZA). The day prior to infection, 1.4×10^4 target cells were seeded into the wells of µ-Slide 8 Well Glass Bottom slides (Ibidi). At the time of infection cells were incubated with virus for two to four hours (depending on the time frame of 3D live cell imaging) followed by substitution with the respective culture medium.

### Vectors, lentivirus production and nucleofection

SunTag expression plasmids pHRdSV40-NLS-dCas9-24xGCN4_v4-NLS-P2A-BFP-dWPRE and pHR-scFv-GCN4-sfGFP-GB1-NLS-dWPRE described in the original publication ^8^ were a gift Ron Vale (Addgene #60910 and # 60906). Due to low lentiviral titers of the dCas9 construct, which are most likely due to the size of the plasmid. We removed four copies of the GCN4 peptide sequence using XhoI sites, that were already present in the vector, and replaced the BFP fluorescence protein by puromycin resistence using present NotI sites and standard cloning methods.

IFI16, KDM2B, SP100, LaminB and H2B were cloned as C- or N-terminal mCherry fusion constructs into LeGO-vectors^17^ for stable transgene expression. To generate a KDM2B zinc finger mutant (KDM2BmutZF), we replaced the KQ motif, that directly interacts with unmethylated cytosine residues within the KDM2B zinc finger ^44^, as well as the two subsequent cysteine residues, which are important for the formation of the zinc finger structure, by glycine.

For the stable expression of optimized sgRNAs, we first subcloned the U6 promoter BbsI sgRNA cloning/expression cassette from the pSpCas9(BB)-2A-GFP (PX458) CRISPR editing vector that was a gift from Feng Zhang (Addgene #48138)^45^ into pCR2.1 (Invitrogen). Then, we replaced the stem loop sequence of the sgRNA by the enhanced extended stem loop sequence and A-U flip described by Chen and colleagues ^15^. gRNAs TR1, TR2, TR3 and TR4 were then inserted into the pCR2.1-U6-chen according to the protocols protocols of the Zhang lab. After confirming correct sgRNA sequences by sanger sequencing, we cloned the entire expression cassette into a Zeocin resistance LeGO vector ^17^ (http://www.lentigo-vectors.de) for generation of lentiviral particles.

For the generation of the SunSeT vector we used the backbone (including the LTRs) of the pCyL50 piggyBac vector (Sanger repository, Wellcome Trust Sange Institute). By PCR and Gibson assembly we inserted a CAG promoter upstream of the SunTag dCas9-GCN4(24x)-puroR sequence followed by three individual sgRNA expression cassettes and the scFv-sfGFP expression cassette.

The piggyBac transposase expression construct was obtained by the sanger repository (Wellcome Trust Sanger Institute).

Nucleofection of SunSeT and transposase vectors was performed using an Amaxa nucleofector (Lonza Bioscience) and the Ingenio Electroporation Kit for Lonza-Amaxa NucleofectorII devices (MoBiTec: MIR50118).

### Immunofluorescence analysis

For detection of LANA expression we performed standard PFA fixation based immunofluorescence (IF) analysis using rabbit anti-LANA antibody ^23^ and goat anti rabbit Alexa-555 secondary antibody (Invitrogen).

### Microscopy and 3D live cell image acquisition

3D confocal live cell imaging was performed using a Nikon Ti2 confocal spinning disc microscope. Laser intensities and exposure times (300 ms max.) were adjusted prior to imaging to achieve optimal imaging conditions over longer time periods. All images were generated using a 100x oil objective. Images were generated as confocal Z-stacks (0.5 µm step size with 5-10 µm total distance in Z). Time series were acquired using build in functions of the NIS-Nikon Elements AR software v4.5 (Nikon).

### Image processing

Acquired mages were registered to correct signal shifting due to different emission wavelengths if necessary. Only linear adjustments were applied to images to enhance visualization. 3D reconstructions and live cell imaging movies were generated using NIS-Nikon Elements AR software (v4.5) and Imaris X64 8.2.1 (Bitplane). Quantification of SunTag/SunSeT signals was performed with Imaris or Arivis (Zeiss) as indicated.

### ChIP-qPCR and ChIP-seq

ChIP and respective qPCR and primers as well as sequencing and ChIP-Seq data analysis for global enrichment of KDM2B on viral genomes in comparison to host loci have been performed essentially as described earlier ^30^.

## Acknowledgements

We thank the members of the Grundhoff lab for discussion and technical advice. We thank the LIV technology platforms *Microscopy and Image Analysis* and *High Throughput Sequencing* for technical support. This work was supported by the Deutsche Forschungsgemeinschaft (DFG, German Research Foundation) in the framework of the Research Unit FOR5200 DEEP-DV (443644894) through project grants GR 3318/5-1 and BO 4158/5-1 awarded to A.G. and J.B.B., respectively.

## Supplementary data

**Figure S1:**
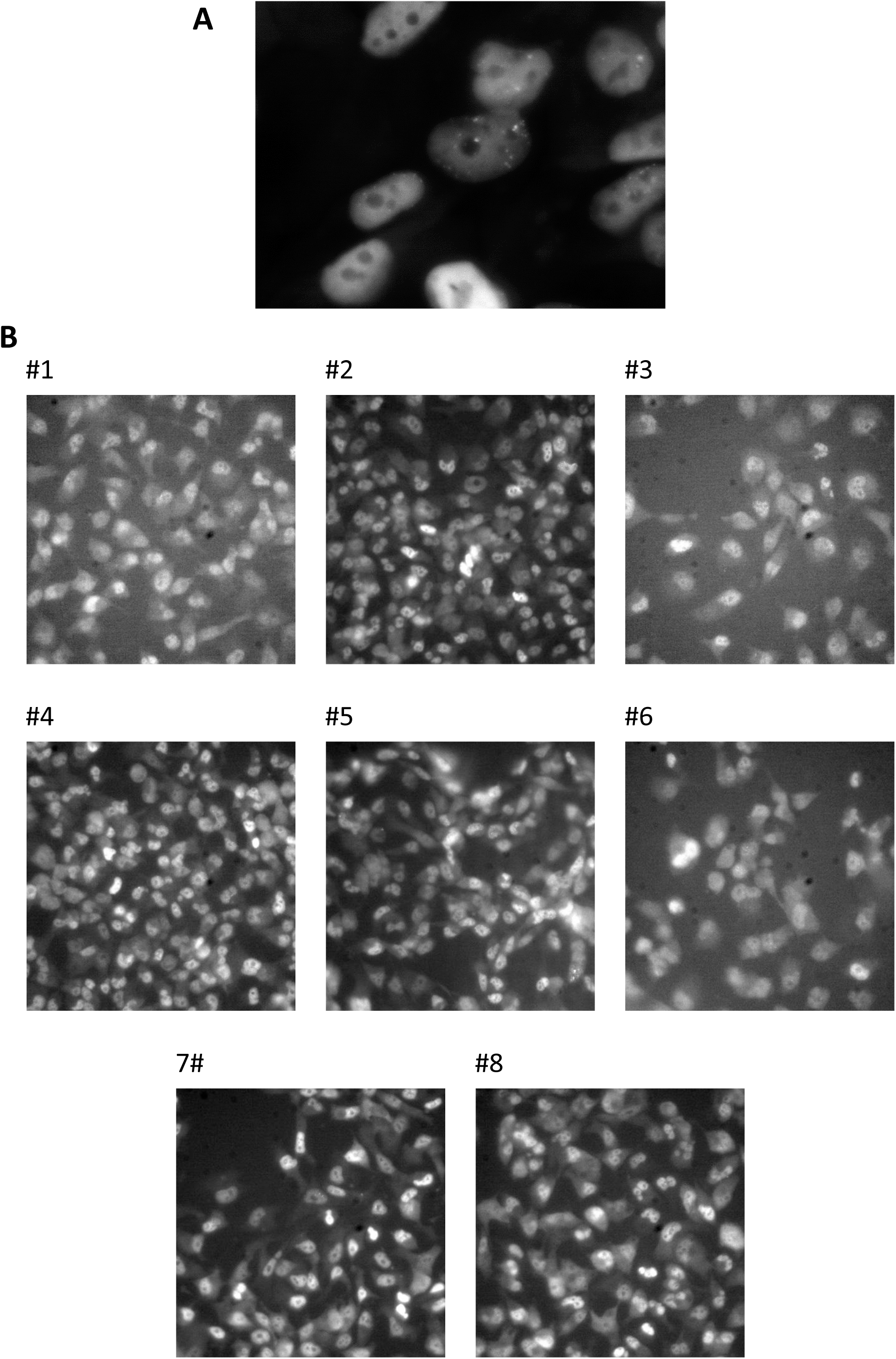
Selection of clones for SunTag KSHV imaging. A) SLK-ST3 bulk cells were infected with KSHV and imaged on a Biostation (Nikon). Newly appearing dot like structures indicated KSHV episomes. B) 46 single cell clones were generated from SLK-TR3, infected with KSHV and screened for best potential visualization of KSHV DNA at 48h p.i. The 8 clones showing the best (GFP) showed dot like structures after infection and imaging by conventional bright field fluorescence microscopy are shown in the micrographs.

**Figure S2:**
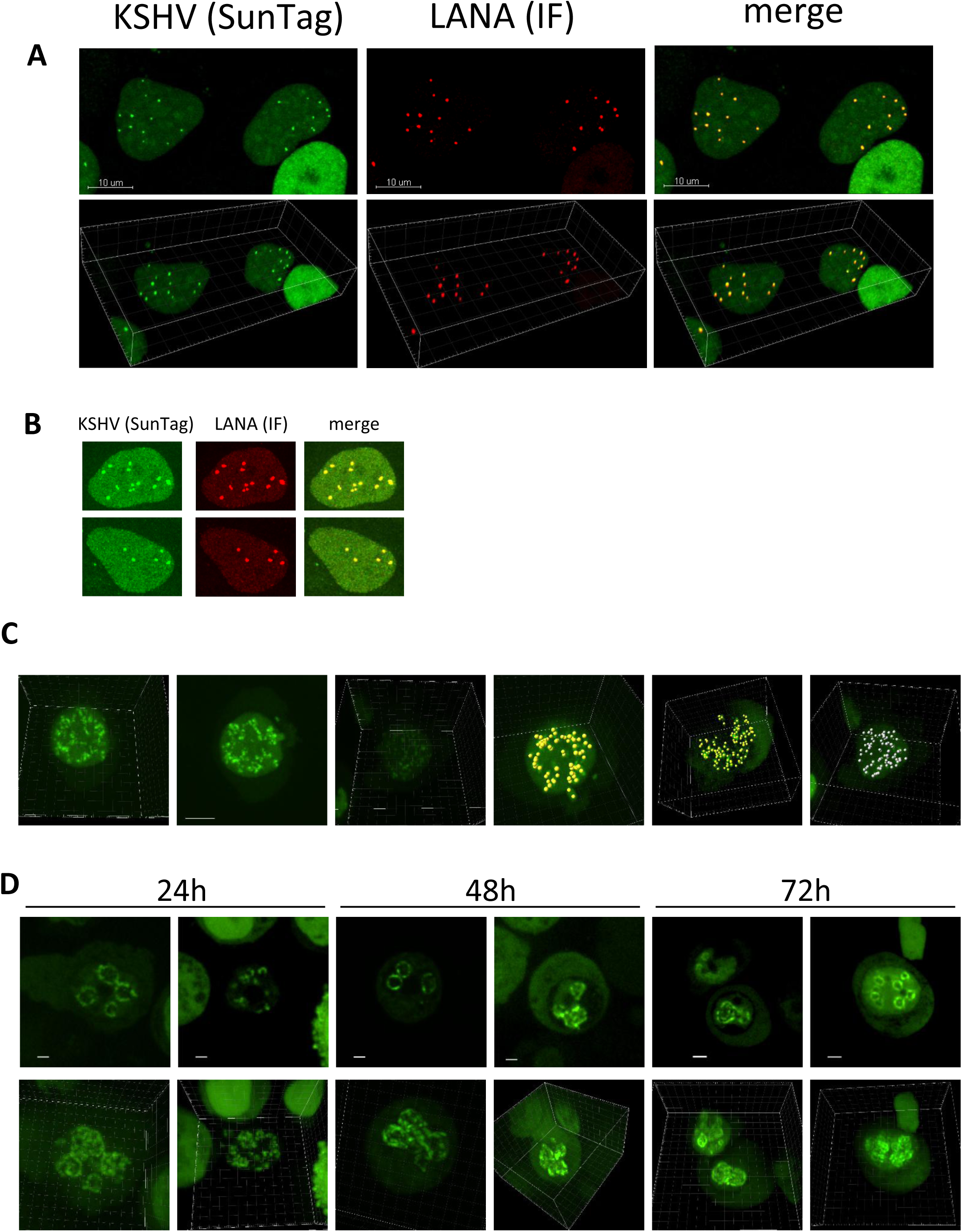
Additional SunTag KSHV images. A) Immunofluorescence of LANA at 48 hours post infection of SLK-ST3#3 with KSHV. Images represent MIP (upper panel) and 3D volume rendering (lower panel). B) Two additional cells from the experiment shown in Figure 1B. C) Three additional representative SunTag KSHV BCBL1 cells (first three images) and examples of dot quantifications by Imaris (yellow or white spheres in the right three images). D) SunTag KSHV BCBL1 cells were lytically induced by TPA/Na+Butyrate treatment and images were acquired at indicated time points post induction. The upper panel shows confocal images and the lower panel 3D reconstruction from Z-stacks.

Movie S1: 3D reconstruction of LANA IF of SLK-ST3 cells at 48 ours post KSHV infection.

Movie S2: 3D reconstruction of BCBL1-ST3.

Movie S3: 3D reconstruction of BCBL1-ST3 at 72 hours post induction with TPA/Na+butyrate.

Movie S4: 3D reconstruction of BCBL1-ST3 at 72 hours post induction with TPA/Na+butyrate.

Movie S5: 3D reconstruction of BCBL1-ST3 at 72 hours post induction with TPA/Na+butyrate.

Movie S6: 3D reconstruction of a BCBL1-ST3 cell transduced with H2B-mCh during mitosis.

Movie S7: 3D live cell imaging of SLK-ST3 cells during the first 4 hours of infection with KSHV (large).

Movie S8: 3D live cell imaging of SLK-ST3 cells during the first 4 hours of infection with KSHV (small).

Movie S9: 3D live cell imaging of SLK-ST3 cells undergoing mitosis.

Movie S10: 3D reconstruction of a SLK-ST3 cell transduced with LaminB-mCh at 8 hours post infection with KSHV.

Movie S11: Z-stack slices of an SLK-ST3 cell transduced with LaminB-mCh at 8 hours post infection with KSHV and calculated colocalization of SunTag and LaminB given as white channel.

Movie S12: Long-term KSHV infected SLK-SunSeT cells transduced with H2B-mCh imaged over 5days

Movie S13: 3D live cell imaging of an SLK-ST3 cell transduced with SP100-mCh from 4 to 12 hours post infection with KSHV.

Movie S14: 3D reconstruction of SLK-ST3 cells transduced with SP100-mCh at 30 hours post infection with KSHV.

Movie S15: 3D live cell imaging of an SLK-ST3 cell transduced with KDM2B-mCh during the first 4 hours of infection with KSHV (overlay).

Movie S16: 3D live cell imaging of an SLK-ST3 cell transduced with KDM2B-mCh during the first 4 hours of infection with KSHV (green; SunTag).

Movie S17: 3D live cell imaging of an SLK-ST3 cell transduced with KDM2B-mCh during the first 4 hours of infection with KSHV (red; KDM2B-mCh).

Movie S18: 3D reconstruction of an SLK-ST3 cell transduced with KDM2B-mCh at 28 hours post infection with KSHV (overlay).

## Notes

### Competing Interest Statement

The authors have declared no competing interest.

